# Spectral waveform analysis dissociates human cortical alpha rhythms

**DOI:** 10.1101/2024.03.16.585296

**Authors:** Janet Giehl, Markus Siegel

## Abstract

The non-sinusoidal waveform of neuronal oscillations reflects the physiological properties of underlying circuit interactions and may serve as an informative biomarker of healthy and diseased human brain function. However, little is known about which brain rhythms can be dissociated based on their waveform and methods to comprehensively characterize waveforms are missing. Here, we introduce a novel spectral waveform analysis (SWA) that provides a complete waveform description, is noise-resistant, and allows to reconstruct time-domain waveforms. We applied this framework to human magnetoencephalography (MEG) recordings during rest and identified several distinct and previously unknown cortical alpha waveforms that were temporally stable and specific for individual subjects. Our findings suggest at least four distinct alpha rhythms in human sensorimotor, occipital, temporal, and parietal cortex. SWA provides a powerful new framework to characterize the waveform of neural oscillations in the healthy and diseased human brain.

## Introduction

Traditionally, neural oscillations have been conceptualized as sinusoidal processes that are characterized by their base frequency and amplitude. However, recent evidence from human EEG, MEG and ECoG recordings suggests that non-sinusoidal waveforms of neural oscillations are highly prevalent in the human brain (Giehl et al., 2021; Lozano-Soldevilla et al., 2016; Schaworonkow and Voytek, 2021). The specific waveform of a neural oscillation reflects the physiological properties of the underlying neural circuit interactions. Thus, waveforms may differ between distinct brain areas, networks and -oscillations, due to genetic variation, in relation to brain disorders, and may change dynamically due to task demands (Cole and Voytek, 2017). Indeed, recent evidence has linked changes of the shape of cortical oscillations to brain disorders, such as Parkinson’s disease (Cole et al., 2017; Jackson et al., 2019) and schizophrenia (Bartz et al., 2019). Together, these findings point to the non-sinusoidal waveform of neural oscillations as a promising new biomarker. Beyond the frequency and amplitude of oscillations, this biomarker may allow to further dissociate and characterize neural oscillations. Furthermore, it may provide a novel window into the underlying physiological processes in the healthy and diseased human brain.

Despite this intriguing potential, to date, the waveform of neuronal oscillations has only been studied sparsely (Cole and Voytek, 2017). Take cortical alpha oscillations as an example. The most well-known example of a characteristic oscillatory waveform in the human cortex is the so called “mu” rhythm (Gastaut et al., 1952; Pineda, 2005; Tiihonen et al., 1989). This arch-shaped alpha-band rhythm is observed over sensorimotor cortex, and its prominent waveform distinguishes it from the more sinusoidal occipital alpha rhythm (Kuhlman, 1978). Together with a third temporal “tau” rhythm, there are at least three distinct alpha oscillations in the human cortex (Klimesch, 1999; Lehtelä et al., 1997; Niedermeyer, 1991, 1990; Tenke and Kayser, 2005). However, the actual number of distinct alpha oscillations is currently still unknown. Furthermore, the three well-described alpha oscillations have originally been distinguished based on their distinct responses to different tasks (Feshchenko et al., 2001; Klimesch, 1999; Tenke and Kayser, 2005). It remains unclear if these functionally distinct rhythms can all be dissociated based on their waveform.

The limited understanding of oscillatory waveforms is also a consequence of methodological limitations. Currently available waveform analysis methods for neuronal data assess only a pre-selected number of waveform characteristics that may not to capture all potentially relevant waveform features (Cole and Voytek, 2017). In addition, waveforms are typically assessed in the time domain, which inherently limits the signal-to-noise ratio (Bartz et al., 2019). In sum, little is known about the potential of non-sinusoidal waveforms to dissociate and characterize human brain rhythms, which is also due to methodological limitations.

To address this, here, we developed a novel spectral waveform analysis (SWA) that comprehensively quantifies waveforms and that is highly resistant to noise. The method assesses waveforms in the frequency domain, by exploiting the inherent relationship that exists between the observable waveform in the time domain and the cross-frequency patterns that define a periodic non-sinusoidal waveform in the frequency domain. We applied this method to human magnetoencephalography (MEG), to investigate if, and potentially how many, alpha oscillations can be dissociated based on their waveform in the human brain. We identified several distinct and previously unknown cortical alpha waveforms that were temporally stable and specific for individual subjects. Our findings suggest at least four distinct alpha rhythms in human sensorimotor, occipital, temporal, and parietal cortex.

## Results

### Spectral Waveform Analysis (SWA)

A non-sinusoidal oscillation (Figure 1B) is defined by its fundamental frequency *f*_1_, amplitude, and waveform. The waveform shape defines the periodic pattern of the oscillation and includes additional information beyond the fundamental frequency and amplitude (Bartz et al., 2019). A non-sinusoidal waveform in the time-domain can be defined as the sum of harmonic sinusoidal components (Figure 1C) that are located at integer multiples of the waveform’s fundamental frequency (Figure 1A). The fundamental frequency of the waveform alone merely represents a sinusoidal signal (top sinusoid in Figure 1C), whereas the entire waveform can be reconstructed as the sum over all sinusoidal harmonic components. Each of the harmonic components has a specific phase-shift *ϕ*_*k*_ and amplitude *A*_*k*_ relative to the fundamental component. Jointly with the fundamental frequency, these phase-shifts and relative amplitudes comprehensively define the waveform. Formally, the spectral representation of a periodic waveform can, thus, be defined in the form of a discrete Fourier series (Figure 1, bottom).

**Figure 1.**
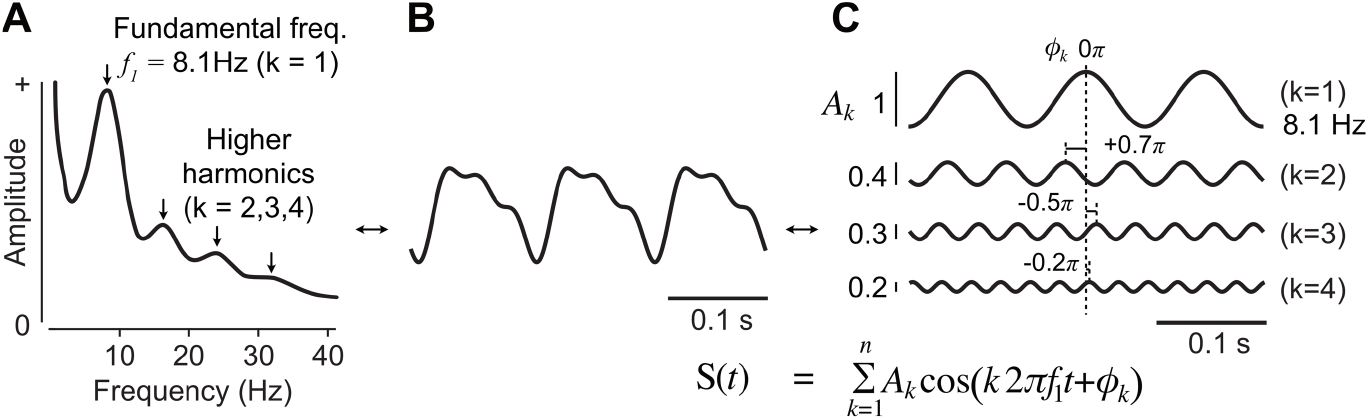
Fourier-series representation of a waveform. (**A**) Harmonic peaks in the amplitude spectrum (**B**) example non-sinusoidal waveform (**C**) harmonic sinusoids that, in their sum, reconstruct the shape of the waveform in (B). (bottom) The formal representation of this sum is a Fourier series. The fundamental frequency *f*_1_, the relative amplitudes *A*_*k*_ and relative phases *ϕ*_*k*_ of all harmonic sinusoids jointly encode the waveform information.

To derive the relative phases and relative amplitudes of harmonic components, our approach exploits a characteristic property of non-sinusoidal periodic waveforms: For a stationary waveform, there is a stable cross-frequency phase- and amplitude relationship between the fundamental frequency and its harmonics. Thus, the relative phases and amplitudes of harmonics that represent the waveform are the relative phases and amplitudes of those signal components that have a stable cross-frequency phase relationship to the fundamental frequency. The bispectrum specifically captures these signals with stable phase relationships between different frequencies and disregards signal components without such a relationship (Sigl and Chamoun, 1994). Thus, the bispectrum is particularly noise resistant (Bartz et al., 2019; Giehl et al., 2021). Furthermore, the bispectrum allows to derive the relative phases and amplitudes of exactly those signal components that are phase-coupled to the fundamental frequency and characterize the waveforms of interest (see Methods and Appendices A and B). This renders the bispectrum, and its amplitude-normalized form, the bicoherence, particularly well-suited for waveform analysis. Based on these insights, we devised a spectral waveform analysis (SWA) that involves two steps. First, the fundamental waveform frequency *f*_1_ is located by determining the presence and spectral location of harmonic bicoherence peaks. Harmonic bicoherence peaks indicate a stable phase-relationship across harmonic frequencies and are, thus, a sensitive measure of oscillations with non-sinusoidal waveforms (Bartz et al., 2019; Giehl et al., 2021; Lozano-Soldevilla et al., 2016). Second, the relative phases *ϕ*_*k*_ and amplitudes *A*_*k*_ of harmonics are derived from the bispectrum. The relative phases *ϕ*_*k*_ can be iteratively reconstructed from the bicoherence phase estimates between successive harmonics (see Methods and Appendix A). The relative harmonic amplitudes *A*_*k*_ can be derived from the bispectrum by applying a spectrally specific normalization factor (see Methods and Appendix B).

SWA effectively translates the relevant bispectrum estimates into the Fourier-series waveform parameters. This approach has critical advantages. First, SWA only evaluates those harmonic signal components that have a stable phase relationship to the fundamental frequency. Therefore, SWA estimates are particularly resistant to spectrally overlapping noise or other signals that do not show a stable cross-frequency relationship. This substantially increases the noise-resistance compared to time-domain approaches (Bartz et al., 2019). Second, SWA provides a complete waveform description. Thus, it does not require the predefinition of waveform parameters. Furthermore, the complete description allows to reconstruct time-domain waveforms from the estimated frequency-domain parameters.

### Cortical peaks of alpha waveform stability

We applied SWA to human MEG data to investigate if, and potentially how many, alpha rhythms can be dissociated based on their waveform. We analyzed resting-state MEG data of 89 subjects (2 sessions per subject) from the human connectome project (HCP) (Van Essen et al., 2013). We employed linear beamforming (Van Veen et al., 1997) to directly investigate neural activity in cortical source space taking into account the phase-ambiguity of the source-reconstructed signals (see Methods). If functionally distinct cortical alpha networks were associated with distinct alpha waveforms, these cortical alpha networks could be located as distinct spatial peaks of alpha waveform stability. To localize such regions of interest (ROIs) with peaked waveform stability, we mapped the first harmonic bicoherence peak in the alpha-frequency-range (Figure 2A) across the cortex (Figure 2B).

**Figure 2.**
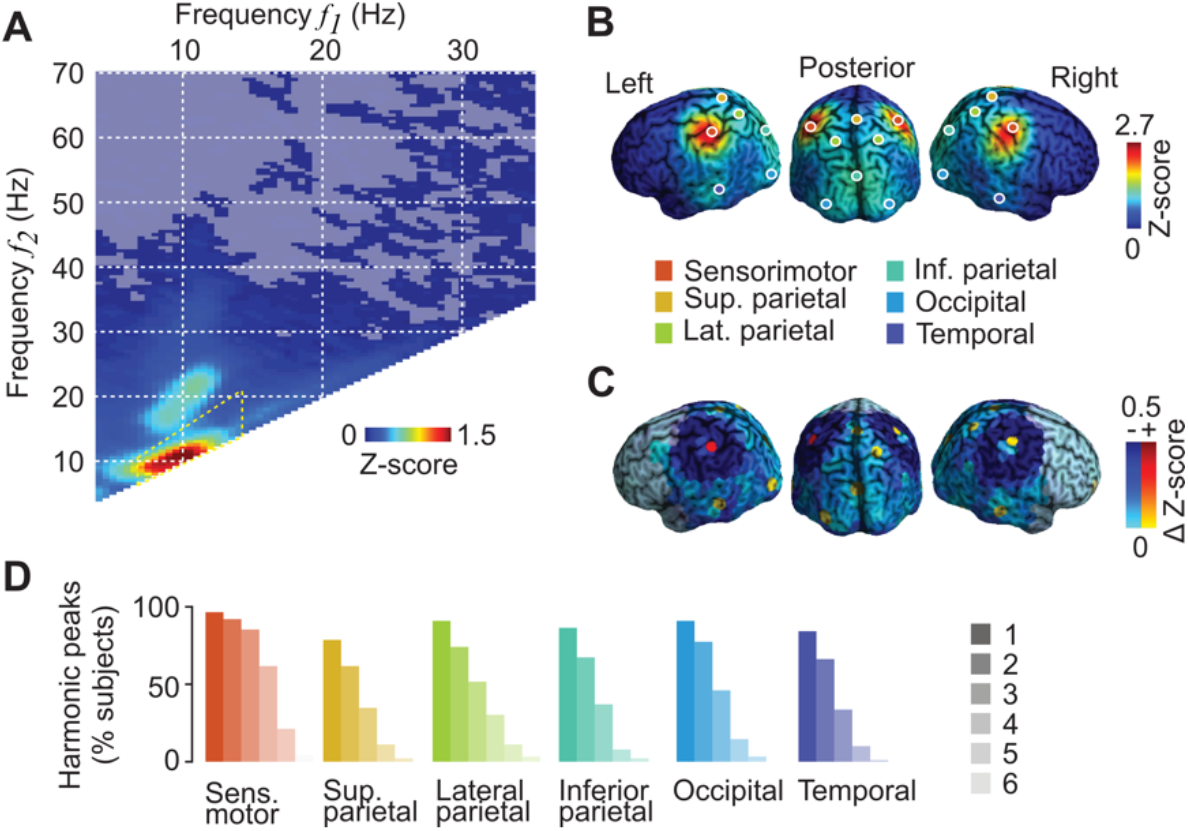
Cortical regions of interest. (**A**) Bicoherence averaged across the brain and subjects. Bicoherence is z-scored against the Null-distribution estimated using circularly time-shifted surrogates. The alpha-frequency range around the first alpha-harmonic bicoherence peak that was used in (B) and (C) is marked in yellow. Non-significant bicoherences (p > 0.05 corrected) are masked. (**B**) ROI locations and cortical distribution of bicoherence averaged across the frequency range marked in (A) and subjects (**C**) Difference between the local and maximum neighbouring bicoherence z-score; local z-scores < 0.5 are masked. Values larger than 0 indicate local bicoherence peaks that were used to define ROI locations. (**D**) Percentage of subjects that showed up to 6 consecutive significant harmonic bicoherence peaks at the different ROIs, which corresponds to the number of coupled higher harmonics of alpha.

This revealed a highly structured pattern of bicoherence (Figure 2B) with several local cortical maxima (Figure 2C). We found lateralized peaks in sensorimotor, parietal, temporal, and occipital regions, as well as medially in inferior and superior parietal cortex. In sensorimotor and temporal cortex bicoherence showed symmetric bilateral peaks. In occipital and lateral parietal areas bicoherence peaked unilaterally, in the left and right hemisphere, respectively. For these unilateral peaks, we included the homologue locations in the contralateral hemisphere, which showed very similar bicoherence (<0.05 difference of z-scored bicoherence between hemispheres). In total, we identified 6 regions of interest (4 bilateral and 2 medial) that showed peak alpha waveform stability, and for which we subsequently analyzed alpha waveforms.

For every subject, cortical ROI, and session, we identified the fundamental alpha frequency based on the corresponding bicoherence spectrum. Then, we identified up to 6 consecutive alpha harmonic peaks with significant bicoherence (p < 0.05, corrected) and extracted the corresponding waveform parameters (relative phase and amplitude). We found two or more significant harmonic bicoherence peaks, that is, three or more harmonic components, in more than half of the subjects at each of the 6 cortical regions (Figure 2D). We identified at least three or four significant harmonic peaks in the lateral parietal and sensorimotor regions, in half of the subjects and at least one hemisphere. We found the lowest (34 %) and highest (85 %) percentage of subjects with three significant harmonic peaks in the temporal and sensorimotor regions, respectively. The percentage of subjects with four or more harmonic peaks ranged from 8 % in the inferior parietal to 62 % in the sensorimotor region. Due to the low percentage of subjects with four or more harmonic peaks, we statistically compared waveforms including the first three harmonics, i.e. we included waveform parameters for harmonic frequencies up to 4*f*_*α*_.

### Distinct alpha waveforms

If the identified peaks of alpha bicoherence reflected distinct alpha rhythms with distinct underlying circuit interactions, then the waveforms of corresponding alpha-oscillations may differ between the corresponding cortical regions. Thus, we next used the derived SWA parameters (fundamental frequency, relative harmonic amplitude and phase) to test for waveform differences between regions (Figure 3). We employed a permutation-based within-subject MANOVA that combined all waveform parameters into one test. Importantly, the overall strength of harmonic coupling as measured by the absolute bicoherence was not included in the test, as differences in this parameter may reflect differences in SNR, rather than a change of waveform shape. Furthermore, we ensured that the pattern of bicoherence strength across harmonics and the pattern of relative amplitudes were compatible with harmonic coupling (see Methods) (Giehl et al., 2021).

**Figure 3.**
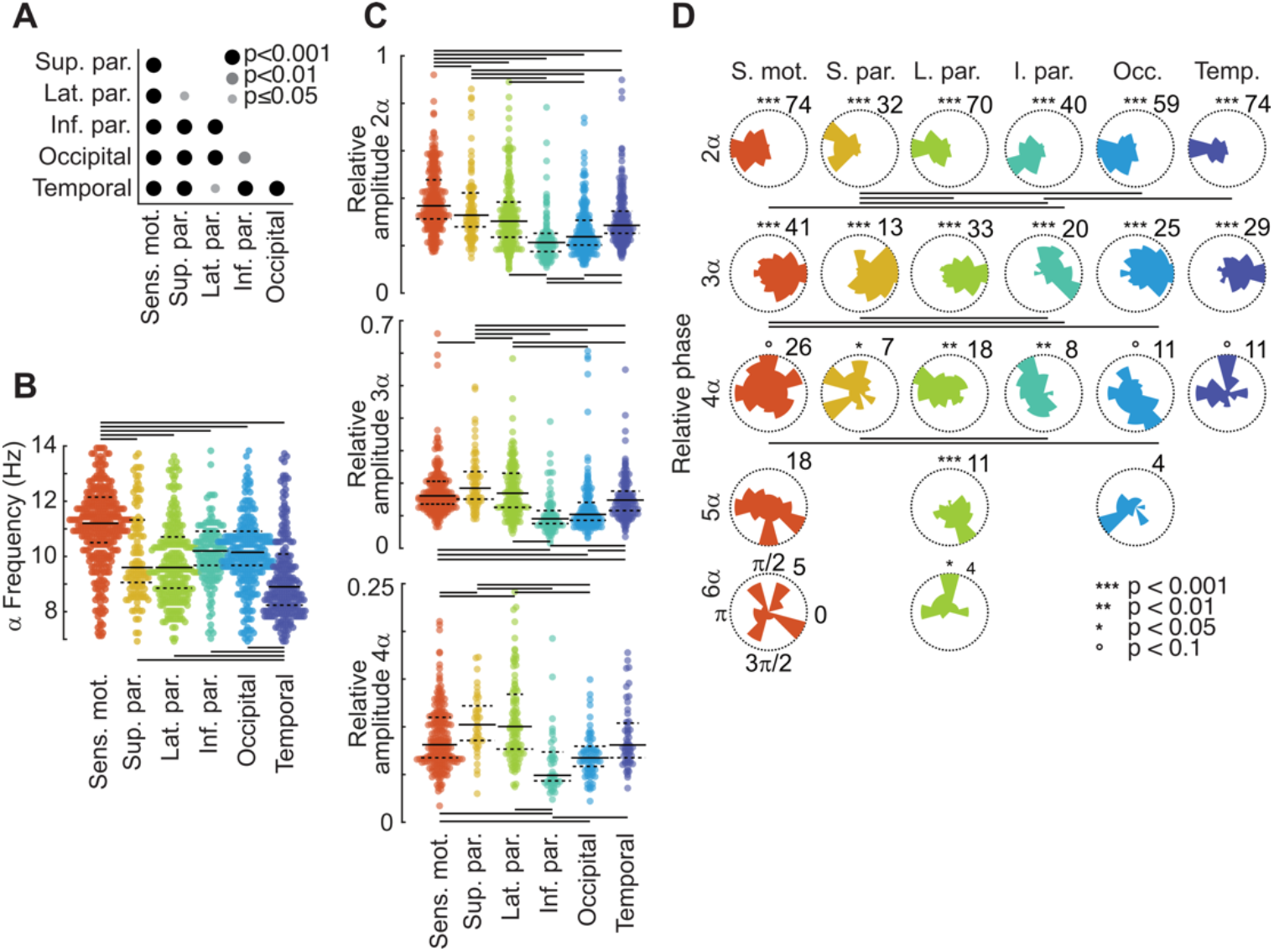
Waveform parameters. (**A**) Significance of pairwise waveform differences between cortical regions (corrected p-values) (**B**) Distributions of bicoherence alpha peak frequency. Dots correspond to individual subjects, sessions and hemispheres. Solid and dashed lines mark median and interquartile range, respectively. Solid lines on top and bottom mark significant differences between regions (p < 0.05; corrected). (**C**) Distributions of relative harmonic amplitude of the first three higher harmonics of alpha. Same display as in (B). Long solid lines mark significant differences between regions (p < 0.05; corrected). (**D**) Histograms of reconstructed relative harmonic phase, for all higher harmonics with at least 15 single observations. Only the first three relative harmonic phases were used for statistical comparisons between regions. Asterisks indicate significant phase non-uniformity (corrected for multiple comparisons). The numbers at the top right of the circular distributions indicate the radial axis scale.

As the MEG data was recorded in two separate 6 min sessions, we first tested for waveform differences between the two recording sessions. We found no significant differences between sessions for any of the 10 cortical regions (all p > 0.05, uncorrected). Similarly, for the 4 bilateral regions there was no significant difference of the waveform between hemispheres (all p > 0.05, uncorrected). We next tested for wave-form differences between the 6 cortical regions. This revealed a highly significant difference between regions (p < 0.001). We followed up with pairwise tests between all 6 regions (Figure 3A). Waveforms significantly differed for all 15 region pairs (all p < 0.05, corrected). Thus, SWA uncovered significant waveform differences between 6 cortical regions. Importantly, we performed the statistics as random-effects analyses on the population level. Thus, the significant waveform differences between regions, in turn, imply the consistency of each region’s waveform across individuals.

We next performed post-hoc tests (permutation-based ANOVAs, see Methods) to investigate, which specific parameter differences were underlying the distinct waveforms (Figure 3B, C and D, see Methods). For the fundamental alpha-frequency, we found that in sensorimotor cortex it was significantly higher, and in temporal cortex it was significantly lower than in all other regions (all p < 0.05, corrected). The fundamental frequency in superior and lateral parietal regions was marginally lower than in inferior parietal and occipital regions (p < 0.1, corrected). Also, the relative amplitude (Figure 3C) and relative phase (Figure 3D) of higher harmonics showed significant differences between several pairs of cortical regions (p < 0.05, corrected). The sensorimotor region had the strongest second harmonic, while the third and fourth harmonic were strongest in lateral parietal cortex. Overall, the inferior parietal region had the lowest relative amplitude for all 3 higher harmonics, and waveforms were most similar between occipital and inferior parietal cortex, which only differed in the relative amplitude of the second harmonic. In general, the number of significant differences decreased for higher harmonics, which is expected as the amplitude, and thus SNR, of higher harmonics decreased (Figure 3C).

As indirectly inferred above, all parameters indeed showed a consistent clustering across individual subjects. For example, the lateral parietal region showed a consistent phase clustering across subjects even up to the sixth harmonic (p < 0.05). However, despite this consistency on the population level, it should be noted that all parameters showed a substantial variability across individuals (Figure 3B, C and D).

In summary, SWA allowed us to dissociate 6 cortical alpha waveform shapes: bilateral sensorimotor, parietal, occipital and temporal waveforms, as well as medial superior and inferior parietal waveforms.

### Alpha waveform stability

The previous analyses revealed a substantial variability of waveform parameters across individuals. Does this variability merely reflect measurement noise or the subject specificity of waveforms? In the latter case, the waveform parameters of each subject should be reproducible across the two recording sessions. To test this, we correlated waveform parameters between sessions across subjects (Figure 4). Almost all waveform parameters were significantly correlated across sessions (p < 0.05, corrected). The lowest and highest waveform stability in terms of cross-session correlation was generally found in occipital and lateral parietal regions, respectively. The fundamental alpha frequency was significantly correlated across sessions for all regions (all p < 0.05, corrected). As for the regional specificity above, cross-session correlation generally decreased for higher harmonics as expected for decreasing SNR. In sum, we found that alpha waveform parameters were correlated between two separate recording sessions across subjects. In turn, this implied that the variability of waveform parameters across subjects was not merely attributable to noise, but reflected subject-specific waveforms that were stable across time.

**Figure 4.**
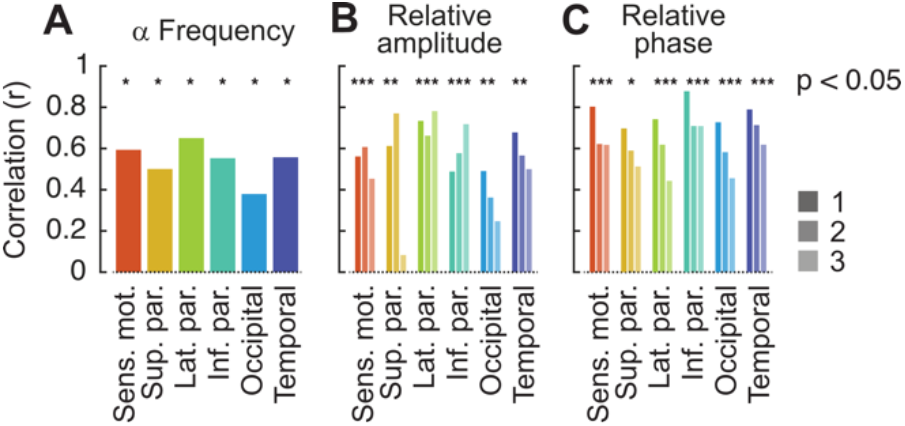
Waveform stability. (**A**) Correlation of fundamental alpha frequencies across two recordings sessions. (**B**) Correlation of relative harmonic amplitudes across recordings sessions. (**C**) Correlation of relative harmonic phases across recordings sessions. Shadings denote harmonic order. Asterisks mark significance (p < 0.05; corrected).

### Reconstruction of alpha waveforms

SWA provides a complete description of oscillatory waveforms. Thus, SWA allows to reconstruct time-domain waveforms from the derived spectral waveform parameters. We applied this approach to visualize waveforms in the time domain, to compare reconstructed waveforms to continuous raw data and to quantify time-domain features of the reconstructed waveforms (Figure 5).

**Figure 5.**
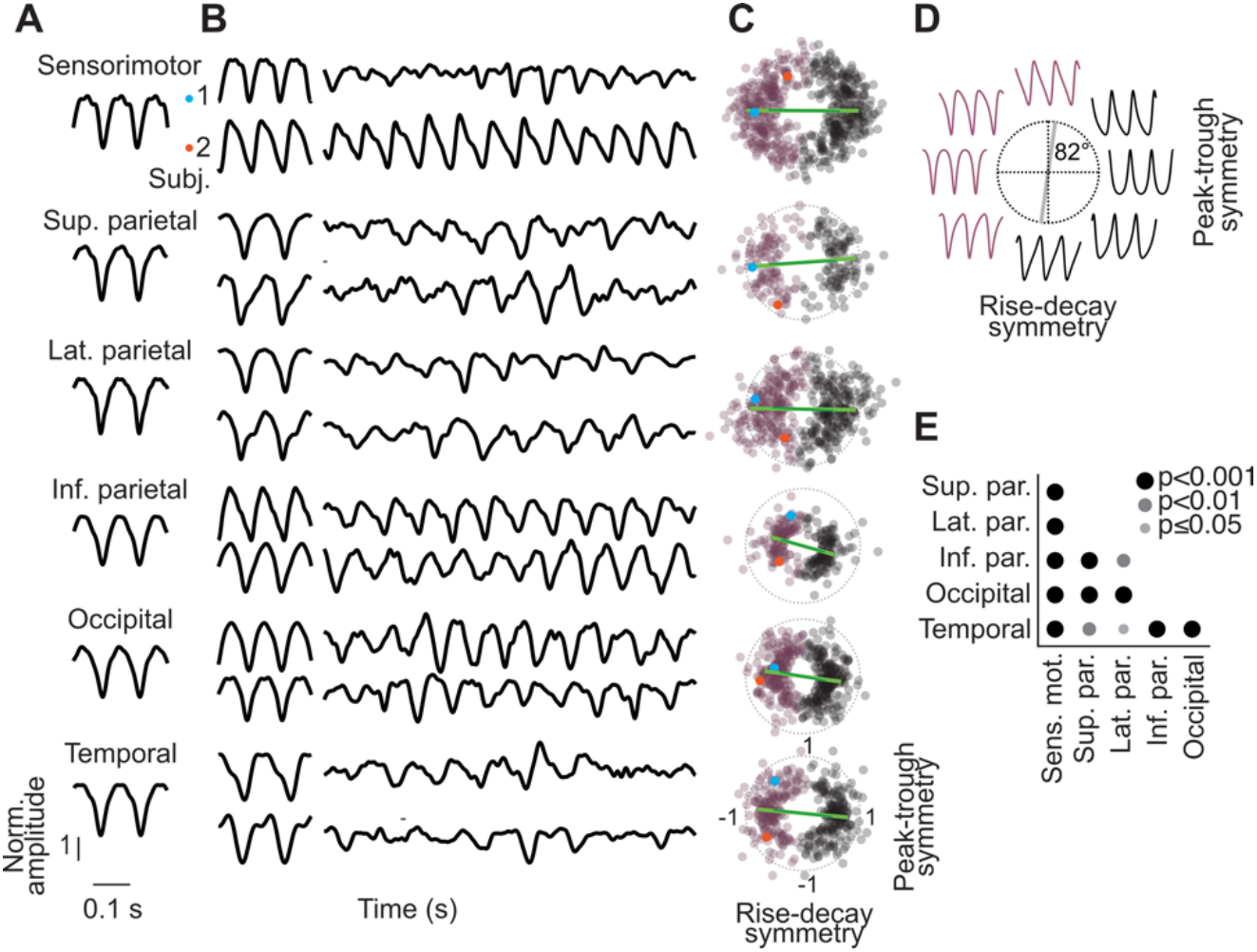
Waveform reconstruction. (**A**) Reconstructed population-average alpha waveforms for all cortical regions of interest. (**B**) Reconstructed wave-forms and raw data of two exemplary subjects (subject 1: top; subject 2: bottom). (left) Reconstructed alpha waveforms (right) Excerpts of raw source-level activity centred around the time with maximum alpha amplitude, low-pass filtered at 55Hz. (**C**) Distribution of rise-decay and peak-trough symmetry of all reconstructed waveforms (violet) and of their amplitude-inverted form (black; 180° phase-flip). Both wave-form polarities are shown to account for the phase-am-biguity of the source-reconstructed MEG signal (see Methods). Dots are individual subjects, sessions and hemispheres. The example subjects shown in (B) are marked in blue (subject 1) and red (subject 2). Dark green lines indicate the circular average of symmetries across subjects. Light green lines indicate the symme-tries of the reconstructed population-average wave-form shown in (A). (**D**) Schematic illustration of waveform shapes with different peak-trough and rise-decay symmetry. The distribution of all single-subject, single-region waveforms was minimal at the indicated 82° axis (compare also panel C). Thus, to account for the 180° phase-ambiguity of source-reconstructed data waveforms to the right of this axis (black circles in C) were amplitude inverted. (**E**) Significance of pairwise waveform differences based on peak-frequency, rise-decay symmetry and peak-trough symmetry (corrected p-values).

For each cortical region, we reconstructed the population average alpha-waveform by averaging all waveform parameters across subjects before projecting and merging the harmonics in the time-domain (Figure 5A). The population average waveforms showed clear non-sinusoidal features. For example, the sensorimotor waveform, displayed the typical “mu”-rhythm waveform that included secondary peaks and troughs nested on top of the fundamental 11.2 Hz crests. Averaging waveform features across subjects may effectively reduce region-specific waveforms shape that are variable but present on the single subject level. To investigate this, we also reconstructed time-domain waveforms of individual subjects. Indeed, the reconstructed individual waveforms (Figure 5B left, two example subjects) did not only show marked variability between subjects, but also suggested more pronounced differences between regions than the population average waveforms.

SWA characterizes waveforms by focusing on harmonic signal components that are phase-coupled across all provided data. This raises the question to what extend the extracted waveforms can even be identified in ongoing neural activity. To descriptively investigate this, we plotted the cortical time courses, centered on the time of maximum alpha amplitude for the same two example subjects and all regions (Figure 5B right). Despite substantial variability of the ongoing activity, for all regions, the reconstructed waveforms resembled at least some of the oscillatory cycles of ongoing cortical activity. This suggested, that SWA effectively extracted the characteristic oscillatory waveforms from ongoing activity.

### SWA-based rise-decay and peak-trough symmetry

SWA-based waveform reconstruction allows to apply time-domain analysis on the reconstructed waveform. Two useful time-domain waveform parameters are the peaktrough- and rise-decay symmetries. Together, these two features capture the global non-sinusoidal shape of a waveform (Figure 5D). To investigate if these time-domain features captured the regional specificity of the SWA-derived waveform, we computed the peak-trough- and rise-decay symmetries of all reconstructed waveforms. To account for the phase-ambiguity of the source-reconstructed data, we performed the analysis on the original and amplitude-inverted waveforms (Figure 5C, violet and black dots) and identified the minimum of the joint distribution across all waveforms at 82° (Fig. 5D). We the performed the following analysis on the waveforms to the left of this axis (Figure 5C, violet dots; see Methods). We statistically compared the waveform symmetry parameters between regions as originally done for the spectral parameters. Indeed, also the time-domain parameters of the reconstructed waveforms captured their regional specificity (Figure 5D). While waveform differences were generally less significant than for the original SWA parameters (compare Figure 3A), almost all pair-wise comparisons between regions yielded significant waveform differences (p < 0.05, corrected; Figure 5E). Again, there were no significant effects of session or hemisphere on the waveforms (all p > 0.05, uncorrected). Thus, SWA-reconstructed waveforms can be successfully used to characterize and dissociate waveforms in the time domain.

## Discussion

Here, we introduce spectral waveform analysis (SWA) – a new framework to characterize the waveform of neural oscillations based on their harmonic profile in the frequency domain. We applied SWA to human resting-state MEG and found that several distinct cortical alpha waveforms can be detected and distinguished. Besides occipital, sensorimotor (mu) and temporal (tau) alpha rhythms, we found evidence for additional parietal alpha rhythms with a distinct waveform. Cortical alpha-waveforms were stable across recording sessions, had a characteristic non-sinusoidal profile across the population, and showed substantial subject-specific variability.

### Analyzing waveform shape in the frequency domain

SWA characterizes oscillatory waveforms in the frequency-domain and critically extends previous time-domain approaches for characterizing wave shapes. First, SWA is particularly noise resistant. The key reason for this robustness is that, by using the bispectrum, the algorithm does not only consider individual frequencies but the relationship between frequencies. Different to the more familiar power spectrum, the bispectrum specifically characterizes stable phase dependencies between frequencies, while disregarding unrelated signals (Sigl and Chamoun, 1994). As these stable phase dependencies between harmonic frequencies are the defining feature of non-sinusoidal waveforms, the bispectrum is ideally suited to extract the waveform information in the frequency domain. Indeed, bicoherence, which is the normalized bispectrum, was already shown to be particularly noise resistant (Bartz et al., 2019; Giehl et al., 2021) and to outperform time-domain waveform analysis when oscillatory signals may not even be directly observable in the time domain (Bartz et al., 2019). Thus, SWA allows for studying waveforms of comparatively weak neural oscillations, which may be particularly advantageous for non-invasive investigations using EEG or MEG. Indeed, the present results uncovered previously unknown alpha waveforms in the human brain using MEG, which highlights the potential of SWA to non-invasively characterize human brain rhythms.

Second, SWA provides a complete waveform description that encompasses all waveform information that can be extracted from the data. The comprehensiveness of this approach has several advantages. First, SWA is not limited to a set of pre-defined waveform parameters that may potentially miss relevant information. Second, and notwithstanding this, SWA allows to reconstruct time-domain waveforms and to then apply well-established wave shape parameters such as e.g., peak-trough and rise-decay symmetry. This approach may be particularly useful as it effectively combines the noise-resistance of SWA with a projection of the high-dimensional spectral parameters space to a lower dimensional, potentially more interpretable parameter space in the time-domain. Finally, because of its comprehensiveness, SWA provides optimal sensitivity to identify differences and modulations of oscillatory waveforms. Indeed, we found that SWA parameters outperformed peak-trough and rise-decay symmetry, in conjunction with waveform frequency, in terms of their sensitivity to distinguish between different waveforms. Thus, SWA can render even very detailed waveform patterns, that may be missed by time-domain parameterization, accessible to statistical analyses.

### Dissociating multiple alpha rhythms

To date, three different alpha rhythms, i.e., neural oscillations at about 10 Hz, have been recognized in the human brain. These rhythms have been distinguished based on their distinct functional responses and cortical source locations (Feshchenko et al., 2001; Klimesch, 1999; Tenke and Kayser, 2005): A prominent visual alpha that is suppressed during eye opening (Berger, 1929), the idiosyncratic motor mu-rhythm (Gastaut et al., 1952; Pineda, 2005) and a third temporal tau-rhythm, which is generally less observable than the other two. The bilateral occipital, sensorimotor and temporal alpha waveforms that we identified conform well with these previously identified alpha oscillation sources. Our results show that these three rhythms express distinct alpha waveforms that are consistent across individuals but also show subject-specific variability.

### Additional parietal alpha waveforms

Our results uncover three additional parietal alpha-waveforms that were distinct from the three previously identified alpha rhythms. Can these distinct waveforms be interpreted as three distinct parietal alpha rhythms? Several factors need to be considered. Volume conduction induces signal mixing, which may result in intermediary waveforms at locations between the primary sources of original rhythms (Bartz et al., 2019). However, all identified regions of interest marked local maxima of waveform stability, which is difficult to explain as the effect of signal mixing between a smaller number of original rhythms that have a unimodal spatial distribution. However, it should be noted that waveform differences were relatively weak for some pairs of regions. In particular, the waveforms of the middle and lateral parietal regions were quite similar and differed only in their second harmonic phase and third harmonic amplitude. Similarly, occipital and inferior parietal waveforms only differed in the relative amplitude of their second harmonic. These weak differences may well reflect mixing between spatially close but distinct alpha rhythms. However, it cannot be excluded that these weak differences may also reflect spurious intermediary waveforms due to mixing of rhythms with complex multimodal spatial distributions.

The parietal region that most likely showed a previously unknown distinct alpha rhythm was the lateral parietal cortex. The lateral parietal waveform was clearly different from the nearby sensorimotor waveform with both, significantly higher and lower relative harmonic amplitudes. Furthermore, even though the lateral parietal region was located between the inferior parietal and sensorimotor regions, the fundamental alpha frequency was not intermediary to that of the other two regions. Furthermore, the lateral parietal region showed the most consistent relative harmonic phase, which was significantly non-uniform up to the sixth harmonic. Our finding of a distinct parietal alpha rhythm accords well with previous findings suggesting that parietal and occipital alpha oscillations may represent functionally distinct rhythms (Barzegaran et al., 2017; Haegens et al., 2014; Nuttall et al., 2022; Sokoliuk et al., 2019).

Together, our findings suggest that there are at least four different alpha oscillations with distinct waveforms in the human brain. This includes at least one parietal alpha rhythm in addition to sensorimotor (mu), temporal (tau), and occipital alpha rhythms. Future studies are required to delineate if the additional fifth and sixth alpha waveform reflect additional genuine alpha rhythms.

### A new window into neural circuit interactions

SWA opens a new window into neural circuit interactions. The waveform-based dissociation of seemingly monolithic brain rhythms into distinct neural oscillations renders these oscillations accessible as separate biomarkers for functional studies and translational applications. For example, the differentiated evaluation of distinct alpha oscillations may allow to pinpoint their distinct functional roles in the healthy brain or their distinct alterations in brain disorders. Indeed, alpha sub-bands have already been functionally dissociated (Klimesch, 1999, 1997; Klimesch et al., 1998, 1996; Wu et al., 2015) and differentially related to neurodevelopmental disorders (Debnath et al., 2020; Murias et al., 2007; Van der Lubbe et al., 2019). Such alpha sub-band specific effects may well reflect distinct underlying oscillations that could be dissociated based on their waveform.

Furthermore, the waveform itself may be a valuable biomarker. The waveform of an oscillation provides additional information beyond its fundamental frequency and amplitude. Like the fundamental frequency (Donner and Siegel, 2011; Siegel et al., 2012), the waveform of an oscillations is shaped by the biophysical properties and the underlying circuit interactions (Cole and Voytek, 2017; Krishnakumaran et al., 2022). Thus, changes of these circuit interactions due to different functional or cognitive states, neuromodulation (Radetz and Siegel, 2022), or pathological conditions may be reflected by corresponding waveform changes. Indeed, recent findings support this notion and have linked changes in waveform parameters to brain disorders, such as Parkinson’s disease (Cole et al., 2017; Jackson et al., 2019; O’Keeffe et al., 2020) and schizophrenia (Bartz et al., 2019). Unraveling the link between waveform features and underlying circuit mechanisms may allow to infer changes in circuit interactions from changes in waveform. Along the same line, the stable subject-specificity of waveforms that we found in the present study may allow to infer individual differences in circuit mechanisms that may also be linked to genetic variability.

SWA is well applicable to brain rhythms beyond the alpha-frequency range and to invasive recordings. The latter may be particularly useful to relate non-sinusoidal waveforms to underlying circuit mechanisms and spiking activity. Furthermore, SWA can be applied in a time-resolved fashion. SWA is well-suited for a trial-locked approach and could also be considered in a sliding-window setting. This allows to characterize the temporal variability of waveforms, which may provide additional informative biomarkers.

## Conclusion

To conclude, here we introduce a novel spectral waveform analysis (SWA) that characterizes the harmonic structure of oscillatory waveforms. SWA provides a complete waveform description, is noise-resistant, and allows to reconstruct time-domain waveforms. Based on this framework, we identified several distinct and previously unknown cortical alpha waveforms in the human brain. SWA provides a powerful new framework to characterize the waveform of neural oscillations in the healthy and diseased human brain.

## Acknowledgements

We thank Paul Hege for helpful discussions. This research was supported by the European Research Council (ERC) StG 335880 (M.S.) and CoG 864491 (M.S.).

## Author contributions

JG: Conceptualization, Software, Spectral waveform analysis method, Formal analysis, Visualization, Writing – original draft, Writing – Review & Editing

MS: Conceptualization, Supervision, Resources, Project administration, Funding acquisition, Writing – original draft, Writing – Review & Editing

## Data availability statement

The HCP data are available for download from https://www.humanconnectome.org.

## Materials and Methods

### MEG Recording & Preprocessing

We analyzed MEG data from the Human Connectome Project (Van Essen et al., 2013). We used the first two 6-minute resting-state recordings that were available for 89 subjects. During the recording, subjects were in supine position and fixating a red fixation cross on a dark background.

The data were band pass filtered between 0.1 and 400 Hz, and notch filters were applied at 60±1Hz and all higher harmonics. Temporal segments with prominent artefacts were removed as defined in the “baddata” HCP pipeline. Muscle-, eye- and heart-related artefact ICA components were identified by visual inspection and subsequently removed. The set-up and recording is described in detail in (Van Essen et al., 2013).

T1 weighted MRIs were used to warp the individual brain space onto a common MNI source space. We used the corresponding transformation matrices as provided in the HCP data set to warp individual subject space onto a common source model with 457 equally spaced source positions positioned about 0.7 cm beneath the pial surface (Hipp and Siegel, 2015). Source-level activity was estimated using linearly constrained minimum variance (LCMV) beamforming (Van Veen et al., 1997). For each source, the data was projected on the spatial orientation of maximum variance (first spatial eigenvector). This projection induces a 180 º phase ambiguity of the signal because the polarity of the eigenvector direction is arbitrary. We accounted for this ambiguity as detailed below.

We performed a Hanning windowed FFT on 1 s segments to compute bicoherence for the detection of ROIs with alpha waveform stability peaks. Windows were centered at 0.5 s intervals (half overlapping), and zero-padded to 2 s, which resulted in 0.5 Hz binning in the frequency domain. To extract waveform parameters, we then performed Hanning windowed FFT on 1 s segments, centered at 0.125 s intervals. Windows were padded to 10 s, which resulted in 0.1 Hz binning in the frequency domain. Using FFT as described above, we obtained complex estimates in the form of 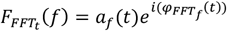, where *a*_*f*_ (*t*) reflects the frequency-specific and time-dependent amplitude at time t and 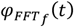 the frequency-specific phase relative to the beginning of each data segment, rather than the phase at time t on which the segment was centered. To account for this discrepancy, we reconstructed the phase at time t for all times and frequencies as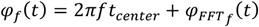, where *t*_*center*_ = 0.5 s, the center of the 1s long Hanning windowed data segments. These phase-corrected estimates 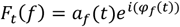 were used for all further analyses.

### Bicoherence

For each recording session, bicoherence *B (f*_*1*_,*f*_*2*_) was computed as:

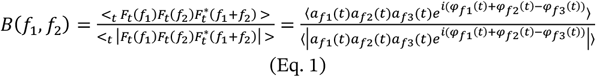

*F*_*t*_*(f)* represents the complex time-frequency transformation of the data at frequency f and time t. <_*t*_ … > indicates the temporal average, which was taken across the entire recording session of 6 minutes,*a*_*f1*_(*t*) is the amplitude time-series at frequency *f*_*1*_, *φ*_*f*1_(*t*) is the corresponding phase time-series, and *f*_3_ =*f*_1_+*f*_2_ The numerator is the bispectrum and the denominator is the normalization factor according to Hagihira et al. (2001). For estimates of coupling strength or “waveform consistency”, we used the absolute value of bicoherence.

For the localization of group-level spatial peaks of bicoherence, we normalized the absolute values of bicoherence to standard Z-scores against the distribution of 100 circularly time-shifted surrogates. To compute the absolute value of bicoherence for a surrogate, the time-series of *f*_3_ was shifted relative to *f*_1_ and *f*_2_.

### ROI selection

To estimate alpha wave-shape consistency on the group level, for each cortical location, we averaged the z-scored bicoherence of the first recording session across the extended alpha frequency range (*f*_1_ from 7 to 14 Hz, inclusive, and *f*_*2*_ up to 1.5 times *f*_1_) and across all subjects. This resulted in a cortical distribution of the coupling strength between the fundamental alpha frequency and its first higher harmonic, which we used as a measure of alpha-frequency wave-shape stability. We identified local maxima of this distribution that exceeded an average z-scored bicoherence of 0.8 and that were not located on the caudal surface of the brain. For the two unilateral non-midline peaks (superior parietal cortex and occipital cortex), we added the homologue location on the other hemisphere.

For each subject, we allowed a small variability of the individual source locations that were used for the respective ROI: we selected the source with the strongest individual alpha-range bicoherence from the average peak location and the adjacent source locations. Adjacent source locations that would also be adjacent to a different ROI were not selected.

### Spectral Waveform Analysis

A periodic waveform *S*(*t*) can be defined as a Fourier series (see also Figure 1):

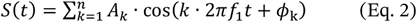

where *f*_1_ is the fundamental frequency of the waveform, *A*_*k*_ represents the amplitude of the phase-coupled *k*-th harmonic relative to the amplitude of the fundamental frequency, with *A*_1_ = 1, and *ϕ*_*k*_ represents the relative phase between the phase of the fundamental frequency and the coupled *k*-th harmonic, with *ϕ*_1_ = 0.

### Fundamental alpha frequency and number of coupled harmonics

To obtain the fundamental alpha frequency *f*_1_, for each subject, ROI and session, we located the individual bicoherence alpha peaks taking into account *f*_*2*_ frequencies from 1 Hz below each *f*_1_ frequency up to 80 Hz. Bicoherence spectra were computed with 0.5 Hz binning. To improve the spectral precision of the detected fundamental and harmonic peak frequencies, we then up-sampled absolute bicoherence spectra to 0.1 Hz binning using spline interpolation. Then, we averaged over *f*_*2*_ and located *f*_1_ with the maximum absolute bicoherence in the extended alpha range from 7 to 14 Hz.

To define the harmonics that were coupled to the identified *f*_1_, we located peaks of absolute bicoherence as a function of *f*_*2*_ for the identified *f*_1_. For each peak, we tested for a significant bicoherence at p < 0.05 using the same time-shifted surrogate statistic employed for the z-scoring of bicoherence and FDR corrected across the entire cross-frequency space (Benjamini and Hochberg, 1995). For a non-sinusoidal oscillation, bicoherence peaks are expected at *f*_*2*_ frequencies that are integer multiples of *f*_1_. Consecutive significant harmonic bicoherence peaks that were located within *f*_*2*_-frequency ranges of *f*_1_ · (*k* ± 0.4) were then defined as the coupled harmonics. The number of significantly coupled harmonics corresponded to the number of these k significant consecutive harmonic peaks.

We ensured that alpha harmonic bicoherence peaks indeed reflected harmonic coupling and could not be due to phase-amplitude coupling between *f*_1_ and 2 * *f*_1_. Harmonic coupling can be confirmed for significant bicoherence coupling peaks, when three or more consecutive harmonic bicoherence peaks have been detected, when the first of two consecutive harmonic bicoherence peaks is not considerably weaker than the second peak, or when only the first harmonic bicoherence peak can be detected (Giehl et al., 2021). Only in 1.8 % of observations (22 cases) the second of two alpha-harmonic bicoherence peaks was numerically stronger than the first. For the remaining 98.2% of observations harmonic coupling was confirmed. The mean and std. of the absolute value of the first bicoherence peak (not z-scored) over all observations was 0.39 and 0.19, respectively, and the mean and std. of the absolute value of the second bicoherence peak was 0.29 and 0.17, respectively. The mean absolute value of the first bicoherence peak of the 22 equivocal observations was 0.16, with a std. of 0.8, for the second bicoherence peak the mean absolute value was 0.19 and std. 0.07. Thus, for these 22 cases, both the coupling and coupling difference were comparatively small. Thus, we included these cases in the further analyses.

### Relative harmonic phase

We reconstructed the pairwise relative harmonic phases *ϕ*_*k*_ from bicoherence phases. We defined *ϕ*_1_ = 0. Relative harmonic phases *ϕ*_*k*_ for k > 1 were reconstructed as the cumulative sum over the harmonic bicoherence phases:

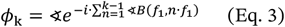

∢ denotes the cosine angle, and ∢B denotes the cosine angle of bicoherence. Eq. 3 exploits a direct mathematical relationship between the phases of harmonic bicoherence estimates and the pairwise relative harmonic phases if harmonic coupling exists (see Appendix A). We used Eq. 3 to obtain the pairwise relative phase between the *k*th harmonic and the fundamental alpha frequency *f*_1_ for all significant higher harmonics.

### Relative harmonic amplitude

To estimate the relative harmonic amplitude *A*_*k*_, we used a normalized bispectrum. We developed a normalization factor that is specifically tailored to retrieve the relative harmonic amplitudes from the bispectrum.

The relative amplitude between the first harmonic (*k* = 1) and itself was defined as)*A*_*f*1_ = 1. Estimates of *A*_*k*_ for higher harmonics (*k* > 1) were reconstructed as:

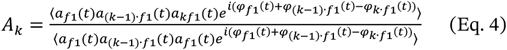

The dividend of Eq. 4 is the bispectrum between a fundamental frequency *f*_1_ and (*k*) 1) *f*_1_. The divisor is the normalization factor. *a*_*k f*1_(*t*) represents the amplitude time course and *φ*_*k f*1_(*t*) the phase time course of the *k*th harmonic frequency and ⟨ ⟩ denotes the average over time. Eq 4 is valid if there is pairwise cross-frequency phase-coupling between all possible pairs of *f*_1_,*f*_*k*™1_, and *f*_*k*_, which is the case for signals with higher harmonics (Appendix B). Theoretically, *A*_*k*_ is real. However, numerically it is usually complex with a phase angle close to zero. Thus, we used the real part of *A*_*k*_ as the measure of relative harmonic amplitude.

To confirm a distribution of ∢ *A*_*k*_ close to 0, we calculated the circular variance *var*_*circ*_ (Z) over the phase angles of all estimates of ∢ *A*_*k*_:

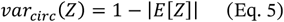

where 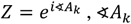 is the cosine phase angle of *ϕ*_*k*_, *E*[] indicates the expected value and | | the absolute value. *var*_*circ*_ (Z) ranges between 0 if all phases *ω* are equal, and 1 for a uniform distribution of phases (Fisher, 1993). Estimates with an absolute phase angle of 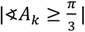 or with a relative amplitude real (*A*_*k*_)) >1, were excluded from further analyses.

### Phase-ambiguity

The 180° phase-ambiguity of the source-reconstructed MEG data can induce spurious differences in the relative harmonic phases between sources. In other words, a random 180º phase-flip of one source with respect to a second source can cause a difference in the relative harmonic phases between these sources. We used a conservative approach to avoid this scenario. Specifically, we aligned all individual wave shapes into one of the two possible directions in such a way to enforce the highest possible similarity between all waveform shapes, given the 180° ambiguity of source-reconstructed MEG data. To this end, we determined which source time-series needed to be flipped by 180º based on the reconstructed waveforms’ peak-trough and rise-decay symmetries. Together, both symmetries span a centrally symmetric space that contains all wave shapes independent of frequency. In this space, a sinusoidal shape is in the center and more asymmetric shapes are more distally. To determine the peak-trough and rise-decay symmetries of waveshapes, we first reconstructed each alpha wave shape *x*(*t*) for every subject and ROI in the time domain using Eq 1 and the derived wave-shape parameters. We reconstructed 5 cycles of each waveform with 1000 equally spaced samples. Peak-trough symmetry) is equal to Pearson’s moment coefficient of skewness 4 (Elgar, 1987) and was computed on *x*(*t*) as:

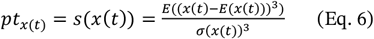

Rise-decay symmetry *rd* is equal to the negative skewness of the imaginary part of the Hilbert transform) (Elgar, 1987) and was computed as:

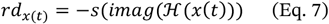

We computed these symmetries for the original and amplitude inverted version of each waveform. In the space spanned by) on the x-axis and *rd* on the y-axis, we located that axis through the origin that intersected the least number of wave shape estimates in the joint distribution of all these wave shape estimates (using an axis width of 0.2 estimated in steps of 1°). This axis was the axis connecting through 82° and 262°. Waveforms with [*pt, rd*] representations located to the bottom-right of this axis were mirrored in amplitude and the waveform parameters of the flipped version was used.

### Distinguishing waveforms

We used multivariate permutation statistics akin to a nonparametric within-subject (repeated measures) MANOVA to statistically assess differences of waveform parameters. Effects were evaluated using 7 dependent variables: the fundamental alpha frequency *f*_1_, the relative harmonic amplitudes *A*_*k*_ and relative harmonic phases *ϕ*_*k*_ for *k ∈* {2,3,4}.

The general outline of the statistical procedure was as follows. We first tested for significant waveform differences across the two sessions, separately for all 10 cortical ROIs. As there was no significant session effect, we kept “session” as an untested factor in the following tests. Next, we tested for significant hemisphere differences, separately for all 4 bilateral ROIs. As there was no significant hemisphere effect, we kept “hemisphere” as an untested factor in the following tests. Then, we performed the main statistical test if waveforms differed across ROIs. This main test was followed up by pairwise post-hoc tests for all ROI-pairs. Finally, second-order post-hoc tests assessed which specific waveform parameters differed between which ROIs.

All nonparametric permutation MANOVAs followed the same logic. For *f*_1_, the three *A*_*k*_ and the three *ϕ*_*k*_ we separately computed F-values, as for a parametric within subject ANOVA. To obtain F-values for the circular relative harmonic phases *ϕ*_*k*_ we transformed all *ϕ* to the form *Z* = *e*^*iϕ*^ (with *ϕ* ∈[−π; π)) and we used multiplication with the complex conjugate in place of squaring to compute circular sums of squares. This way, we obtained 7 separate F-values, one for each of the dependent variables. We then used a permutation statistic to assess the statistical significance against the Null hypothesis of no waveform difference between ROIs. We used 5000 permutations of randomly reassigning ROI-labels. For each permutation we computed 7 permutation F-values. Testing against the distribution of permutation F-values, we obtained one p-value for each of the 7 unpermuted F-values, as well as one permutation p-value for each of the 5000 permutation F-values. We applied Fisher’s method (Mosteller and Fisher, 1948) to pool the 7 independent variables. We derived one X^2^-value from the 7 unpermuted p-values as well as 5000 permutation-X^2^ -values from the permutation p-values:

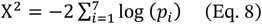

The unpermuted X^2^-value was then tested against the distribution of permutation X^2^-values to obtain the p-value of the null-hypothesis of no difference between ROIs. We applied the same permutation statistic to compute post-hoc MANOVAs for all pairwise ROI comparisons. We corrected the p-values of these pairwise MANOVAs for multiple comparisons using false discovery rate correction (FDR, Benjamini and Hochberg, 1995). For the pairwise ROI-comparisons, the 7 separate F- and p-values of each pairwise MANOVA served as the second-order post-hoc statistic of differences of individual waveform parameters. Again, these p-values were all FDR-corrected for multiple comparisons.

### Cross-session stability

We used Spearman correlation to correlate the fundamental alpha frequency and the relative harmonic amplitudes across the two recording sessions. The relative harmonic phases were correlated using a permutation test, where the phase correlation *r*_*ϕ*_ between the two sessions (here *A* and *B*) was calculated as:

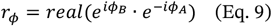

Here, *ϕ*_*A*_ represents the relative harmonic coupling phase of one subject, ROI and harmonic in session). Permutation was performed across subjects. The resulting p-values were FDR corrected for multiple comparisons across the 6 ROIs and 3 higher harmonics (Benjamini and Hochberg, 1995).

### Phase non-uniformities

We tested the non-uniformity of the distributions of the reconstructed relative harmonic phase at every ROI and higher harmonic up to the 6^th^ harmonic. We computed the mean vector length *vl* across all subjects 4 at every ROI and higher harmonic:

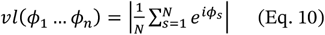

Significance of non-uniformity was determined with respect to the distribution of 1000 *vl*_*U*_(*u*_1_ …*u*_*n*_) drawn from *u*_*s*_ ∼ *u*([0,2π)) The resulting p-values were FDR corrected across all ROIs and harmonics (Benjamini and Hochberg, 1995).

### Group-level waveform reconstruction

To reconstruct the characteristic waveforms on the group level, we averaged the fundamental alpha frequency *f*_1_ and relative harmonic amplitude *A*_*k*_ across subjects. The group-level relative harmonic coupling phases *ϕ*_*k*_ were reconstructed from the group averaged bicoherence phases. Following the results of the tests for phase non-uniformity, subject average waveforms were reconstructed including the waveform parameters from all *k* = 1 … *K* higher harmonics with at least 15 observations and for which phase uniformity of the bicoherence phases was rejected at p < 0.05 after FDR correction. This was *k* = 5 for the sensory-motor and the lateral parietal ROIs, *k* = 4 for the occipital ROI and *k* = 3 for the remaining ROIs. For every ROI, we then applied Eq. 1 to the included group-level waveform parameters.

### Individual waveform reconstruction and temporal excerpts

For each ROI, we reconstructed the alpha for two example subjects using Eq. 1 and the respective waveform parameters. To enable a visual comparison between these Fourier series waveform reconstructions and the original data, we extracted one second of source reconstructed data centered around the time of maximum alpha amplitude for each ROI. The time of maximum alpha amplitude was identified by repeating the time frequency analysis as described above with 10 ms steps and by smoothing the resulting amplitude time series with a 1.5 s Hanning window before locating the temporal maximum. The resulting excerpts were low pass filtered at 55 Hz using a fourth order zero-phase forward-reverse Butterworth filter.

### Peak-trough and rise-decay symmetry

We computed the peak-trough- and rise-decay symmetry for each individually reconstructed waveform *x*(*t*) according to Eq. 6 and Eq. 7 as described in the section “MEG phase ambiguity” above. We tested for waveform differences using the same statistical approach as detailed in section “distinguishing waveforms” above using the three dependent variables alpha frequency *f*_1_, peak-trough symmetry *pt* and rise-decay *rd* symmetry The non-sinusoidal signals *x*(*t*) in Fig. 5d were as:

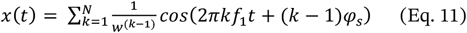

We used *f*_1_ = 10 Hz, *N* = 10, and *φ* _s_ = 0, 0.25π, 0.5π, 0.75π, π, 1.25π, 1.5π and 1.75π for the 8 plotted signals, starting at 0° in counterclockwise order. The waveform constant *w* was defined as the constant fulfilling 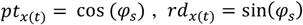 and 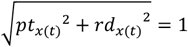 for *N →* ∞ and where ∢ *b*_*k(t)*_ (*f*_1_,*k f*_1_)= − φ _s_for all, *k* ∈ [1, …,*N* −1],, and was numerically approximated as *w* = 2.34521.

### Software

All analyses were performed in MATLAB (MathWorks Inc., Natick, USA) using the Fieldtrip toolbox (Oostenveld et al., 2011) and custom software.

## Appendix

### Appendix A. Reconstructing relative harmonic phase

Eq. 3 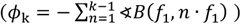 reconstructs the pairwise relative harmonic phases *ϕ*_*k*_ as the negative circular cumulative sum over the corresponding *n* = 1 to *k* = *k*) 1 harmonic bicoherence phases ∢*B* (*f* _1_,*n* · *f* _1_). Eq. 3 is valid, if the bicoherence phase captures the phase-difference between the cross-frequency coherences of two subsequent harmonic waveform components. That is, if:

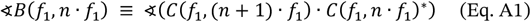

Here, * denotes the complex conjugate,) denotes equivalence, and cross-frequency coherence *C*(*f*_1_, *x* · *f*_1_) is defined as:

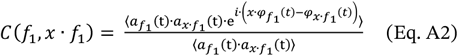

Here, *a*_*f*_(*t*) denotes instantaneous amplitude, *φ*_*f*_(*t*)denotes instantaneous phase, and ⟨ ⟩ denotes the average across time. To conserve a relative simplicity of the notation, we will be omitting the normalization factors in the derivation below.

Eq. A1 can be derived as follows. We momentarily disregard the temporal average and consider only a single point in time *T* with *T ∈ t*. We rewrite the right side of Eq. A1 according to Eq. A2 while omitting the normalization factor (divisor) of Eq A2 because the normalization does not affect phase:

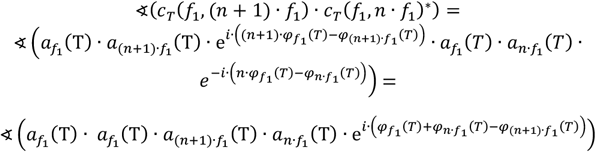

Except for a difference in amplitude weighting, this is identical to the phase of bicoherence ∢*b*_*T*_ (*f* _1_,*n* · *f* _1_), or of its non-normalized form, the bispectrum, considering only a single point in time T with *T∈t*:

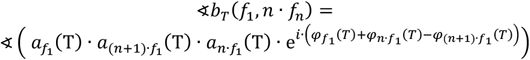

This equivalence remains valid for temporal averages, if the following requirements are fulfilled: there are significant 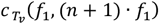 and 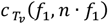 at times *T*_*v*_ (*T*_*v*_ *∈ t*.), and the angle 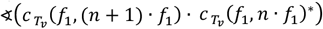 is stationary over *T*_*v*_ . These requirements are fulfilled for harmonic components of stationary, non-sinusoidal periodic waveforms, where all three frequencies (*f*_1,_ *f*_*n*_) and *f*_*n+1*_)are jointly harmonically coupled. Any other contributions to *a*(*t*) and *φ*(*t*) at *f*_1,_ *f*_*n*_ *f*_1,_, or (*n*+1)) 1) · *f*_1_, that are not phase-locked, can be expected to cancel out and, thus, vanish in the temporal average of bicoherence. Thus, it is valid to use Equation 3 to reconstruct relative harmonic phases *ϕ*_*k*_ of non-sinusoidal periodic waveforms.

### Appendix B. Reconstructing relative harmonic amplitude

For a stationary non-sinusoidal periodic waveform, it holds that:

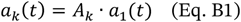

where *A*_*k*_ = *const*. Here, *a*_1_(*t*) is the amplitude time course of the fundamental harmonic; *a*_*k*_(*t*) is the amplitude time course of the *k*th harmonic of the same waveform; and *A*_*k*_ is the stationary relative harmonic amplitude of the *k*th harmonic in relation to the fundamental (*k* = 1) harmonic. We used Eq. 4 to quantify *A*_*k*_:

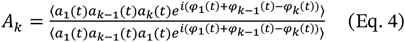

The dividend of Equation 4 is the bispectrum and the divisor is the required normalization factor. Equation 4 can be derived under the assumption of Eq. B1 by substituting Eq. B1 into the right-hand side of Eq. 4:

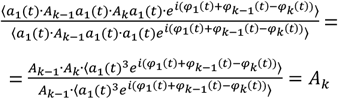

Notably Equation 4 is only valid in the case of harmonic coupling when Eq. B1 can be assumed. In this case, contributions to *a*(t) and *φ*(*t*) that are not phase-locked to the waveform signal can be expected to cancel out, thus, vanish in the temporal average. For practical application, we suggest using the real part of *A*_*k*_ and to exclude estimates with a considerable imaginary component of *A*_*k*_.

